# Rational Designed Hybrid Peptides Show up to a 6-fold Increase in Antimicrobial Activity and Demonstrate Different Ultrastructural Changes as the Parental Peptides measured by BioSAXS

**DOI:** 10.1101/2021.11.05.467440

**Authors:** Kai Hilpert, Jurnorain Gani, Christoph Rumancev, Nathan Simpson, Paula Matilde Lopez-Perez, Vasil M. Garamus, Andreas Robert von Gundlach, Petar Markov, Marco Scocchi, Ralf Mikut, Axel Rosenhahn

**Author notes:** **Correspondence:** Ralf Mikut, Kai Hilpert.

## Abstract

Antimicrobial peptides (AMPs) are a promising class of compounds being developed against multi-drug resistant bacteria. Hybridization has been reported to increase antimicrobial activity. Here, two proline-rich peptides (consP1: VRKPPYLPRPRPRPL-CONH_2_ and Bac5-v291: RWRRPIRRRPIRPPFWR-CONH_2_) were combined with two arginine-isoleucine-rich peptides (optP1: KIILRIRWR-CONH_2_ and optP7: KRRVRWIIW-CONH_2_). Proline-rich antimicrobial peptides (PrAMPs) are known to inhibit the bacterial ribosome, shown also for Bac5-v291, whereas it is hypothesized a “dirty drug” model for the arginine-isoleucine-rich peptides. That hypothesis was underpinned by transmission electron microscopy and biological small-angle X-ray scattering (BioSAXS). The hybrid peptides showed a stronger antimicrobial activity compared to the proline-rich peptides, except when compared to Bac5-v291 against *E. coli*. The increase in activity compared to the arginine-isoleucine-rich peptides was up to 6-fold, however, it was not a general increase but was dependent on the combination of peptides and bacteria. BioSAXS experiments revealed that proline-rich peptides and arginine-isoleucine-rich peptides induce very different ultrastructural changes in *E. coli*, whereas a hybrid peptide (hyP7B5GK) shows changes, different to both parental peptides and the untreated control. These different ultrastructural changes indicated that the mode of action of the parental peptides is different from each other as well as from the hybrid peptide hyP7B5GK. All peptides showed very low haemolytic activity, some of them showed a 100-fold or larger therapeutic window, demonstrating the potential for further drug development.

## 1 Introduction

In the light of the current headlines of the COVID-19 pandemic, the problem of resistance against antibiotics is still progressing and is frequently called a silent pandemic. Especially the beginning of the COVID-19 pandemic demonstrated severely the multitude of effects of an infectious disease where no effective treatment is available. The development of microbial resistance against natural occurring or human-made antibiotics is a natural selection process and was soon discovered after the use of the first antibiotic. With the emergence of multi-drug resistant strains and a dwindling development of antibacterial drugs with new modes of action, the health care systems worldwide are under threat to face a more severe pandemic if no novel antibiotics are developed. Many operations, transplantations, and immunosuppressant therapies may not be performable any longer, leaving modern medicinal care devastated. Hence, it is important and urgent to expand the discovery and translation of new alternatives to treat bacterial infections, especially with new modes of action.

According to the world health organization (WHO, 2019), more than 700,000 people die worldwide each year because of antibiotic-resistant infections. If nothing is done about this antimicrobial-resistant crisis, it was estimated in the O’Neil Report, that the numbers will increase to 10,000,000 by 2050, (https://amr-review.org/Publications.html). Already we are facing extreme multi-drug resistant bacteria like *Mycobacterium tuberculosis, Pseudomonas aeruginosa, Neisseria gonorrhoeae*, and *Staphylococcus aureus*.

Antimicrobial peptides (AMPs) are an extremely diverse group of compounds, found in all kingdoms of life that show various biological functionalities, for example, antibacterial, antifungal, antiviral, antiparasitic, anticancer, and immunomodulatory (Mahlapuu et al., 2016). Several databases are dedicated to AMPs (Kang et al., 2019; Pirtskhalava et al., 2016; Wang et al., 2016). What makes them very interesting for antimicrobial drug development is the fact that they have also numerous modes of action (Brogden, 2005), including inhibiting lipid 2, a cell-wall precursor essential for the bacteria (de Leeuw et al., 2010; Schmitt et al., 2010), blocking the synthesis of important outer membrane proteins (Carlsson et al., 1991), binding to intracellular molecules like histones, RNA, DNA (Cho et al., 2009; Hale and Hancock, 2007; Kobayashi et al., 2000; Xie et al., 2011), DNA-dependent enzymes (Hilpert et al., 2010; Marchand et al., 2006), and ribosomes (Knappe et al., 2016; Krizsan et al., 2014, 2015b; Mardirossian et al., 2014, 2018b, 2019a, 2020). Furthermore, a study investigated the effects of AMP’s on blood components, since most peptides have low bioavailability and therefore the most probable manner of administration is intravenously (Yu et al., 2015). Consequently, the safety and efficacy of some AMPs have been investigated in clinical trials (Czaplewski et al., 2016; Greber and Dawgul, 2017).

Bac5 (43mer) is a proline-rich AMP (PrAMP) from the cathelicidin class, isolated from bovine neutrophils more than 30 years ago (Gennaro et al., 1989). Recently, fragments of Bac5 were investigated and the mode of action was described (Mardirossian et al., 2018a). Bac5 fragments inhibit bacterial protein synthesis as other PrAMPs and are most active against Gram-negative pathogens (Mardirossian et al., 2019b; Tokunaga et al., 2001). The N-terminal 1-17 amino acid fragment of Bac5, called Bac5(1-17), retained antimicrobial activity and the same overall mechanism of action (Mardirossian et al., 2019b). Bac5(1-17) was optimized using spot synthesis and variant 291 (RWRRPIRRRPIRPPFWR-CONH_2_) showed improved antimicrobial activity while displaying low toxicity and the same mode of action as the parental peptide (Mardirossian et al., 2019a).

We have developed a novel prediction method (unpublished results) for AMPs with a low haemolytic activity that was based on our peptide library screen of 3,000 members (Mikut et al., 2016). Two arginine-isoleucine-rich peptides optP1 (KIILRIRWR) and optP7 (KRRVWIIW), were identified from this new prediction strategy, showing broad-spectrum activity. The peptide optP7 was used as a lead and selected for studying lipidation, glycosylation, cyclisation, and grafting into a cyclotide (Grimsey et al., 2020; Koehbach et al., 2021). Hybridization of different AMPs has been investigated to further improve their performance (Shang et al., 2020; Wade et al., 2018, 2019). Here we report the creation of different hybrid molecules using peptides optP1 (KIILRIRWR) and optP7 (KRRVWIIW) in combination with the Bac5(1-17) variant 291 (RWRRPIRRRPIRPPFWR-CONH_2_) and a consensus sequence from proline-rich antimicrobial peptides, called consP1 (VRKPPYLPRPRPRPL-CONH_2_). The antimicrobial activity and haemolytic activity of the parental and hybrid peptides were determined. Small-angle X-ray scattering (SAXS) measurements, performed at the BioSAXS beamline, showed differences in the ultrastructural changes of *E. coli* for two selected hybrid peptides compared to the parental peptides.

## Materials and Methods

### Peptides

Automated solid-phase peptide synthesis (SPPS) on a MultiPep RSI peptide synthesizer (Intavis, Tuebingen; Germany) was used to produce the AMPs. We applied 9-fluorenyl-methoxycarbonyl-tert-butyl (Fmoc/tBu) strategy and used N,N,N′,N′-Tetramethyl-O-(1H-benzotriazole-1-yl)uronium hexafluorophosphate (HBTU) activation. Crude peptides were analysed and purified to the homogeneity of >90% by liquid chromatography-electrospray ionisation mass spectrometry (LC-ESI-MS). For a detailed description please see (Grimsey et al., 2020). The peptide hyP7B5Cys and consP1 were purchased at Synpeptide Co., Ltd (Shanghai, China).

### Bacterial strains

In this project the following bacterial strains were used: a) methicillin-resistant *Staphylococcus aureus* (*S. aureus*) HO 5096 0412 (a neonatal infection isolate, isolated in Ipswich, England in 2005, obtained from Jody Lindsay (St. George’s, University of London), b) *Escherichia coli* (*E. coli*, UB1005, F-, LAM-, gyrA37, relA1, spoT1, metB1, LAMR) used solely for transmission electron microscopy, *E. coli* (ATCC 25922), c) *Pseudomonas aeruginosa* PAO1 obtained from Dr Robert E.W. Hancock (Department of Microbiology and Immunology, University of British Columbia).

### Bacteriological media and culture conditions

To grow the bacterial strains Mueller Hinton broth (MHb) (Merck, Life Science UK Limited, Dorset, UK)) was used, which was prepared concurring to the manufacturer’s directions. Liquid bacterial cultures were incubated at 37°C for 18-20 h on a shaker, bacterial culture on solid media was incubated for 18-24 h at 37°C.

### Minimal Inhibitory Concentration determination

A broth microdilution assay was applied to determine the minimum inhibitory concentrations (MIC) in MHb for both, using 5 × 10^5^ CFU/ml (MICs in Table 1) and 10^8^ CFU/ml (BioSAXS and electron microscopy, values in bracket in Table 1). A detailed description of the method can be found in the open-source publication by von Gundlach et al. (von Gundlach et al., 2019).

**Table 1:**
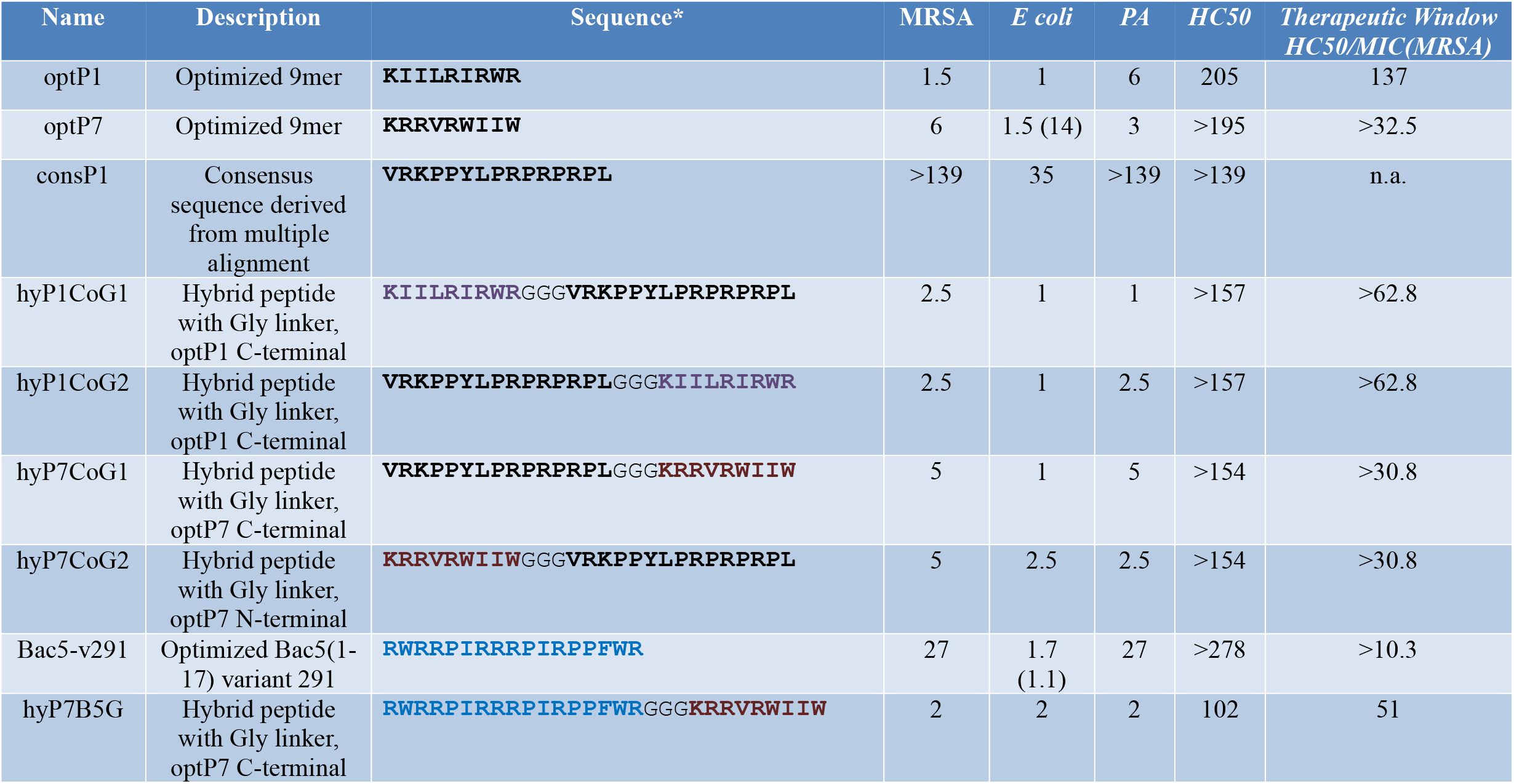
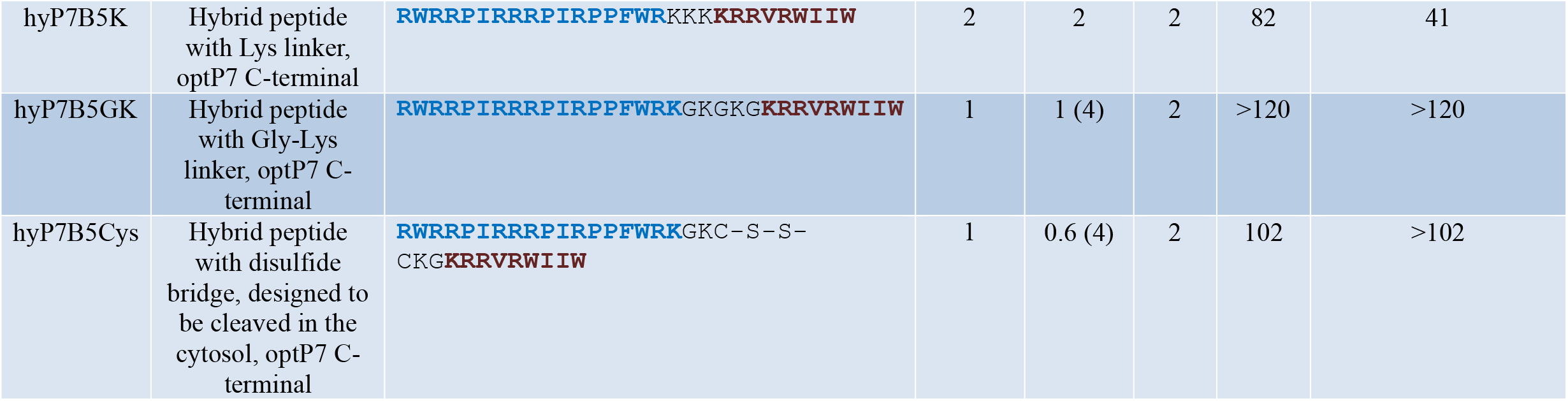
Minimum inhibitory concentrations (MIC) in µM. Data are for n = 3 with modal values reported; all values determined in Mueller– Hinton broth (MHb) using 5×10^5^ CFU/mL bacteria; MRSA = methicillin-resistant *Staphylococcus aureus*, HC50 represents the peptide concentration needed to induce haemolysis in human red blood cells at 50%, values in brackets show MIC values in MH using 1×10^8^ CFU/mL. * All peptides are C-terminal amidated. PA stands for P. aeruginosa, n.a. for not applicable. The therapeutic window is given for MRSA only, as an example for their potential as a drug development candidate.

### Haemolytic activity assessment

The protocol to determine the haemolytic activity of the peptides and HC50 values were described previously (Grimsey et al., 2020). Briefly, human blood was washed and diluted with PBS and incubated with a pre-made dilution series of the peptides and incubated for 1 h at 37 °C. The 100% haemolysis value. Triton X (1% final concentration) was used for Haemoglobin was measured at 414 and 546 nm using an ELISA plate reader. HC_50_ values were determined using the Prism software (GraphPad).

### Sample preparation for BioSAXS, BioSAXS measurement and data evaluation

A detailed description of the sample preparation for BioSAXS, the BioSAXS measurement and consequently the data evaluation methods can be found in the open-source publication by von Gundlach et al. (von Gundlach et al., 2019). In brief, suspensions of the bacteria after different treatments with 2 x MIC_10^8_ (40 min, 37°) were injected by an autosampler (Franke et al., 2012). After multiple exposures, the background was subtracted, and the scattering curves calculated. Data evaluation was done using the toolbox Gait-CAD and SciXMiner (“Peptide Extension”)(Mikut, 2010; Mikut et al., 2017). Data were analysed using a principal component analysis (PCA) using the frequency range 0.05 nm^-1^ to 0.412 nm^-1^.

### Electron microscopy

Sample preparation for transmission electron microscopy was performed by the Image Resource Facility, St Georges University. Overnight cultures of *E. coli* were logarithmically grown to 10^8^ CFU/mL. Untreated and treated with optP7 samples were incubated for specific time points. At the desired time points sample was removed, centrifuged, washed with 0.1 M PIPES buffer pH 7 (Merck Life Science UK Limited, Dorset, UK), and fixed with 2.5% glutaraldehyde in cacodylate buffer (Merck Life Science UK Limited, Dorset, UK). The samples were then washed with PIPES buffer and incubated in 2 % Osmium Tetroxide (Taab Laboratories, Berkshire, UK). A series of ethanol washes followed to dehydrate the samples with 70, 90, and thrice with 100 % ethanol for at least an hour. The samples were then washed with propylene oxide (Scientific Laboratory Supplies Ltd, Nottingham, UK) and transferred as a 1:1 mixture into resin (Agar Scientific, Chelmsford, UK) for 45 minutes. Then this mixture was replaced with 100 % resin and placed at 60°C for 48 hours. Thin sections of resin (80-100nm) were cut using a Reichert-Jung ultracut E Microtome (Leica Microsystems, Wetzlar, Germany) and collected on copper mesh grids (Agar Scientific, Chelmsford, UK). They were then stained with 2% uranyl acetate in an aqueous solution (Taab Laboratories, Berkshire, UK) for 15 minutes followed by Reynolds lead citrate stain (VWR International Ltd, Leicestershire, UK) (Reynolds, 1963) for 2 minutes. Samples were examined using a Hitachi 7100 electron microscope (Hitachi Europe Limited, Berkshire, UK) operating at 75kV.

## Results

Based on the success of the hybridization strategy reported by others, for example (Fox et al., 2012; Shang et al., 2020; Wade et al., 2018, 2019; Xu et al., 2014), we hypothesized that hybrid molecules from two different classes of AMPs, which we have identified in the past as potential antimicrobial drug candidates, may lead to a further improvement of antimicrobial activity. Thus we combined the excellent activity of proline-rich and arginine-isoleucine-rich AMPs. Proline-rich AMPs (PrAMPs), like Bac5, have no lytic mode of action but enter the cytosol by the membrane transporter SbmA and to a small proportion by the MdtM complex (Krizsan et al., 2015a; Mattiuzzo et al., 2007). PrAMPs inhibit protein synthesis by targeting the bacterial ribosome (Krizsan et al., 2014; Mardirossian et al., 2020). The mode of action of the two short (9-13mer) arginine-isoleucine-rich AMPs optP1 and optP7 used in this study is not yet determined. Our current theory is that these two peptides possess a rather “dirty drug” modality by stressing the cell wall by replacing Ca^2+^ ions, depolarizing the cell membrane, enter the cell by passive transport and bind ATP and other negatively-charged molecules like DNA and RNA and hence interact and inhibit various processes in the cell (Hilpert et al., 2006, 2010; Von Gundlach et al., 2016a, 2019). Quite early on in our drug development process, we have selected optP7 over optP1 for a slightly better MIC against *P. aeruginosa*. Further investigations were performed with peptide optP7 only.

Firstly, we investigated *E. coli* cells treated with optP7 by transmission electron microscopy (TEM) to get a better understanding of the mode of action/ultrastructural changes of the peptides on *E. coli*, see Figure 1. In order to achieve a sufficient number of bacteria to form a pellet, a bacterial concentration of 1×10^8^ CFU/mL was used. The MIC of optP7 against *E. coli* at 1×10^8^ CFU/mL was determined at 14 µM, see Table 1. The peptide was incubated with the bacteria for 60 minutes at twice the MIC (28µM). The vast majority of the cells showed no sign of lysis, see Figure 1C. The outer membrane of *E. coli* was damaged and blebbing was visible, where lipopolysaccharides (LPS) form small vesicles and are shed from the cell, see Figure 1. The inner membrane seemed to remain intact. The DNA in untreated bacterial cells comprised dispersed fibrils in the cytoplasm, see Figure 1A. The bacterial DNA is concentrated in a ‘DNA–plasm’ called the nucleoid (Ohniwa et al., 2007). The shape of the nucleoid depends on the ratio of opposing forces (compacting and expanding). Expanding forces are caused by translation, transertion, and translocation of nascent membrane proteins (Cabrera et al., 2009). The dispersed shape in untreated cells by electron microscopy is created by the DNA-dependent RNA polymerase activity. For the ribosome synthesis, this enzyme creates transcription foci and as a consequence twists the nucleoid into its observed shape (Jin et al., 2013). The peptide optP7 creates an enlarged and spherical nucleoid with some small and denser foci, Figure 1 B) and C), typically seen in DNA-dependent RNA polymerase inhibitors and protein synthesis inhibitors. It is in support of our hypothesis that optP7 is able to translocate into the cell and bind negatively charged molecules like ATP, RNA, and DNA and thus inhibit many different important biological activities in the cells (Hilpert et al., 2010). Granular substructures in the cytosol could indicate aggregation of cytosolic proteins. These substructures were observed before using small AMPs (Von Gundlach et al., 2016a, 2019).

**Figure 1:**
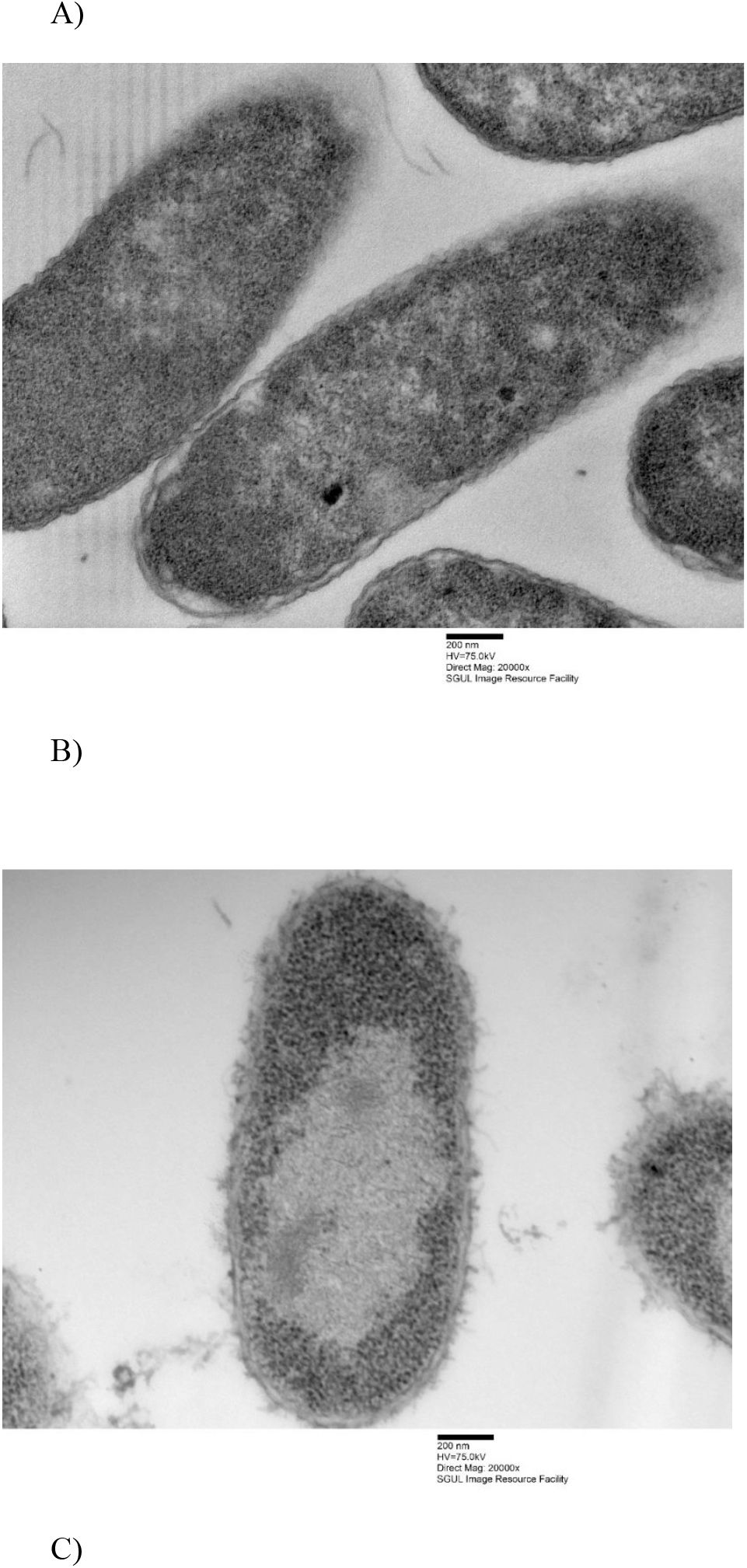
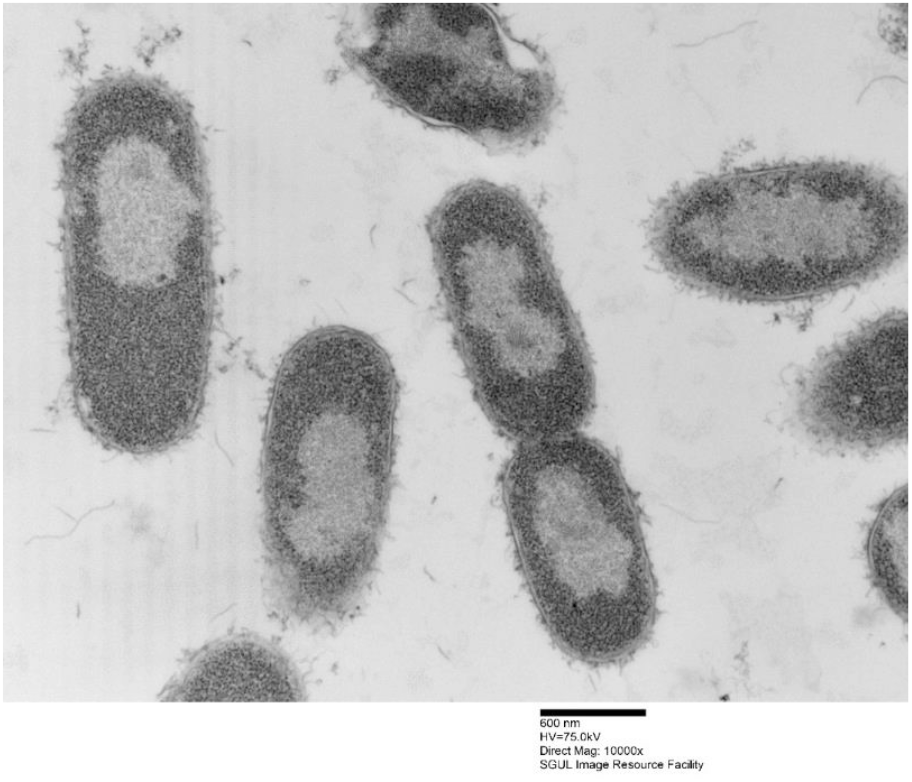
Transmission electron micrographs of *Escherichia coli*, A) untreated samples, B) treated with optP7 for 60 min, 20,000 times magnification, and C) treated with optP7 for 60 min, 10,000 times magnification.

Based on the two arginine-isoleucine-rich peptides optP1 and optP7, hybrid peptides were created by merging with two proline-rich peptides. One PrAMP was the optimized Bac5(1-17) variant 291 (Mardirossian et al., 2019a). The second one was created by a multiple sequence alignment of PrAMPs. From the alignment, a consensus sequence was derived (consP1: VRKPPYLPRPRPRPL-CONH_2_), see Figure 2. Various linkers were tested to combine these peptides, see Table 1.

**Figure 2:**
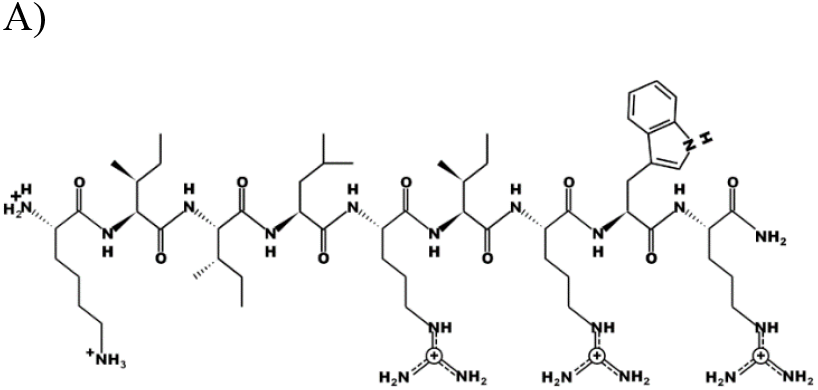
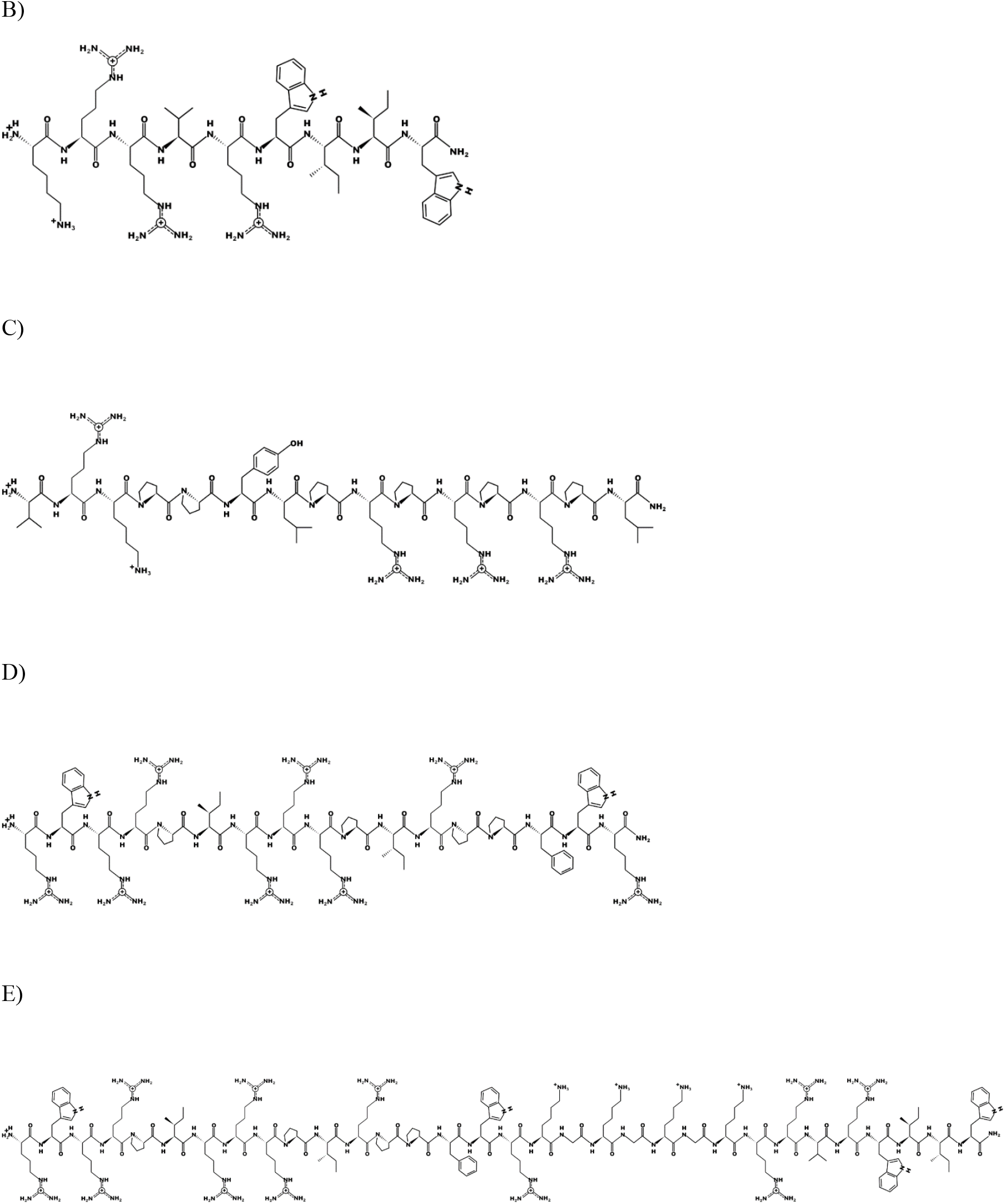
Schematic representation of the parent peptides and one hybrid peptide using the program PEPDRAW (http://pepdraw.com/). A) optP1, KIILRIRWR-CONH_2_, B) optP7, KRRVRWIIW-CONH_2_, C) consP1, VRKPPYLPRPRPRPL-CONH_2_, D) Bac5-v291, RWRRPIRRRPIRPPFWR-CONH_2_, E) hyP7B5GK, RWRRPIRRRPIRPPFWRKGKGKGKRRVRWIIW-CONH_2_

All peptides were tested against the gram-negative bacteria *Pseudomonas aeruginosa* PA01 and *Escherichia coli* UB1005, and the gram-positive methicillin-resistant bacteria *Staphylococcus aureus* HO 5096 0412 (MRSA). The peptides optP1 and optP7 showed strong activity against all tested bacteria, see Table 1. The peptide consP1 showed only activity against *E. coli* and Bac5-v291 showed a medium activity against MRSA and *P. aeruginosa*, and strong activity against *E. coli*. The first four hybrid peptides (hyP1CoG1/G2 and hyP7CoG1/G2) are combinations of optP1 and optP7 with consP1. A flexible glycine linker was used and the influence of the order of peptides within the hybrid molecule was investigated, see Table 1. The combination of consP1 and optP1 resulted in hyP1CoG1 and hyP1CoG2. Both hybrid peptides showed a strongly increased activity against all bacteria when compared to consP1. When compared to optP1 they showed a decrease in activity against MRSA. The antimicrobial activity against *E. coli* remains unchanged in comparison to optP1. In contrast, hyP1CoG1 shows a 6-fold improvement in antimicrobial activity against *P. aeruginosa* and hyP1CoG2 shows a 2.4-fold improvement compared to optP1. The combination consP1 and optP7 resulted in the hybrid peptides of hyP7CoG1 and hyP7CoG2, see Table 1. Both peptides demonstrate a strong improvement in activity when compared to consP1. When compared to optP7, nearby the same antimicrobial activity against MRSA was determined. The peptide hyP7CoG1shows a similar MIC against *E. coli*, hyP7CoG2 shows a 1.7-fold decrease compared to optP7. Activity against *P. aeruginosa* dropped 1.7-fold for hyP7CoG1and was very similar for hyP7CoG2 when compared to optP7.

The next set of peptides was created by combining Bac5-v291 with optP7 using four different linkers. The first was the GGG-linker for flexibility (hyP7B5G), the second a KKK-linker to improve charge (hyP7B5K), the third a GKGKG-linker to combine flexibility and charge (hyP7B5GK). The fourth linker was similar to the third, however, the hybridisation was achieved by a cysteine bridge (hyP7B5Cys). The cysteine bridge is expected to be cleaved in the reducing environment of the bacterial cytosol and both peptides could then act independently once inside the bacterial cytosol.

All the hybrid peptides in this set were 13.5-fold more active against *P. aeruginosa* when compared to Bac5-v291 and showed very similar MIC values when compared to optP7. Peptides hyP7B5G, hyP7B5K, and hyP7B5GK show also very similar MIC values when compared to Bac5-v291 and optP7. The exception is peptide hyP7B5Cys, which showed a 2.5-fold increase in activity compared to optP7 and a 2.8-fold increase compared to Bac5-v291. All the peptides in this set are more active against MRSA when compared to both, Bac5-v291 and optP7. Peptides hyP7B5G, hyP7B5K showed a 13.5-fold improvement, peptides hyP7B5GK and hyP7B5Cys a 27-fold improvement compared to Bac5-v291. Similarly, peptides hyP7B5G, hyP7B5K showed a 3-fold improvement, peptides hyP7B5GK and hyP7B5Cys a 6-fold improvement compared to Bac5-v291. All peptides showed very low haemolytic activity (Table 1). The peptides with the highest haemolytic activity still showed at least a 10-fold therapeutic window, some of them 100-fold or larger.

Small-angle X-ray scattering for biological samples (BioSAXS) on the P12 beamline can be used to measure ultrastructural changes within the bacteria in the range between 2 and 157 nm, from which we used the range from 20 to 120 nm for analysis of the bacteria after treatment with the antimicrobial peptides. In recent years we used this method to classify the mode of action of various antibiotics and antimicrobial peptides against *E. coli* and MRSA (Von Gundlach et al., 2016a, 2016b, 2019). Here we used BioSAXS to investigate the ultrastructural changes of *E. coli* due to the exposition of the selected parental peptides optP7, Bac5-v291, and the hybrid peptides hyP7B5GK and hyP7B5Cys. BioSAXS measurements were performed using bacteria at 1×10^8^ CFU/mL to have high enough bacteria densities for the scattering experiments. The MICs for the selected peptides against *E. coli* at 1×10^8^ CFU/mL were determined (Table 1, values in brackets). BioSAXS measurements were performed at 2x MIC. The corresponding scattering curves of the peptide treated bacteria are presented in Figure 3. To quantify the subtle changes between the different treatments we used a PCA analysis using SciXMiner and the “Peptide Extension” tool (Mikut, 2010; Mikut et al., 2017) (Figure 4). In particular, the two first principal components PC1 and PC2 were used for representation as they provide a suitable and sensitive indicator for structural changes caused by antibiotic treatment (Von Gundlach et al., 2016a, 2016b). The two-dimensional representation of PC1 and PC2 allows to compare the peptide treated bacteria with each other and to the untreated controls. SAXS curves of treated bacteria with similar principal components showed similar internal structures while stronger structural alterations occur if points appear in separated positions in the plot. The proline-rich peptide Bac5-v291 separated very well in the PCA from the short peptide optP7. This shows that the ultrastructural changes induced by these two peptides are very different. This is in agreement with our hypothesis that these two peptides have a different mode of action. The hybrid peptide hyP7B5GK is in between the parent peptides on PC1 and also shows an increase in PC2. The other hybrid peptide hyP7B5Cys is closer to the parent optP7. This hybrid peptide is designed to fall apart in the cytosol due to the reducing environment and it seems that optP7 in this case, drives the ultrastructural changes.

**Figure 3:**
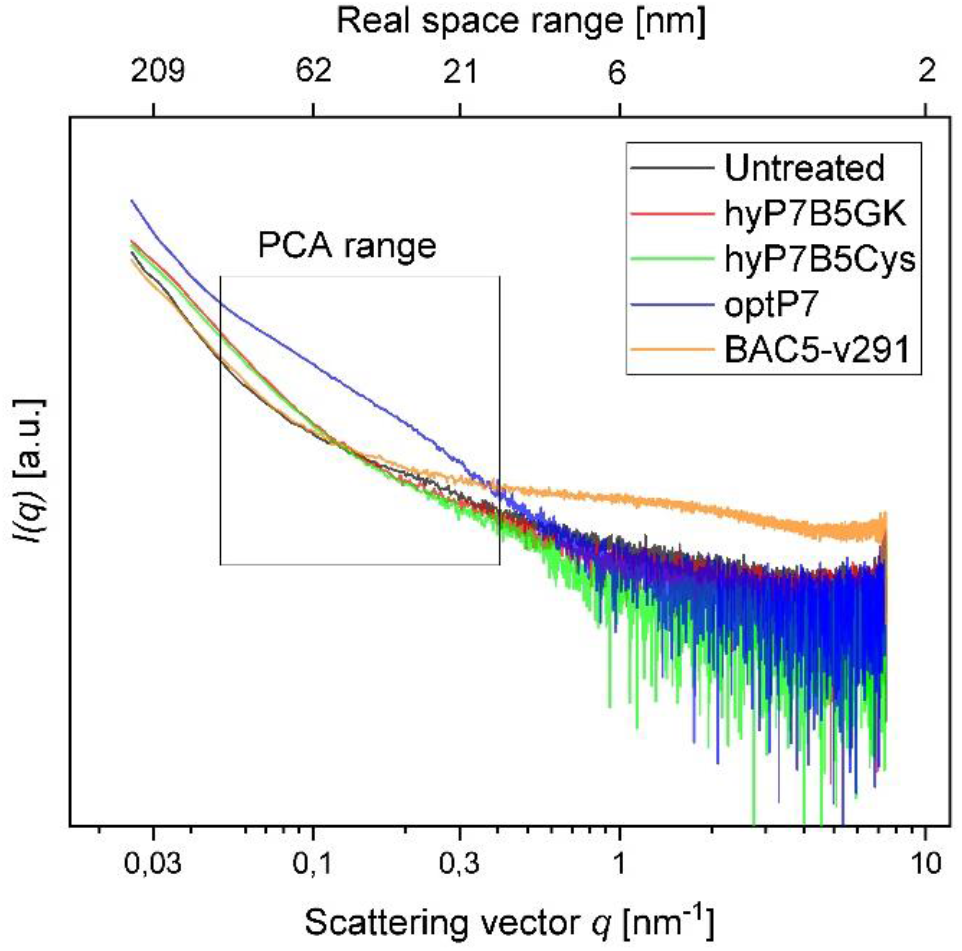
Scattering data from untreated *E. coli*, and treated with four different peptides after 60 minutes, as measured at the P12 BioSAXS beamline at PETRA III (Hamburg, Germany). The principal component analysis (PCA) was calculated using data within the box.

**Figure 4:**
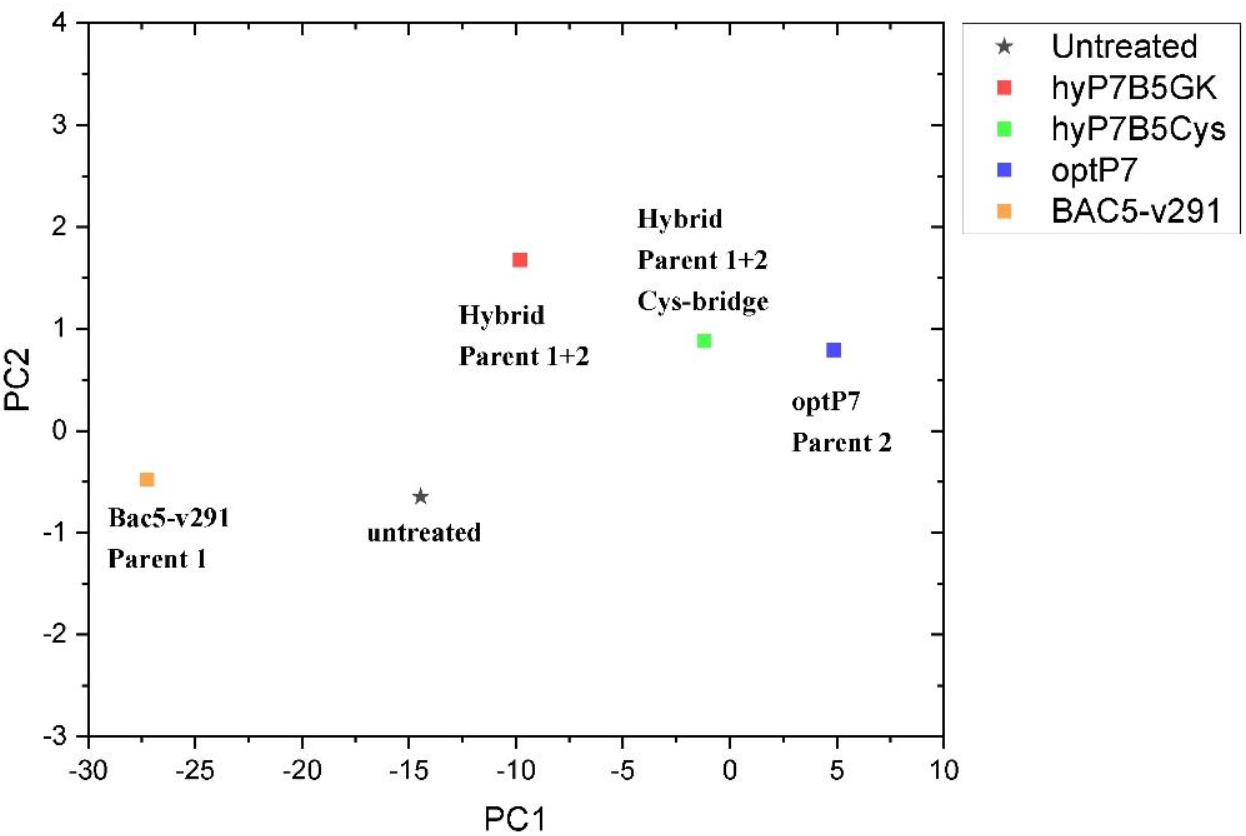
Principal component analysis (PCA) graph, obtained from SAXS curves of *E. coli*, untreated and treated with four different peptides.

## Discussion

Antimicrobial resistance is a silent pandemic that needs urgent solutions in order to be able to provide appropriate health care. Antimicrobial peptides are a very broad and diverse class of compounds that can kill multi-drug resistant bacteria and therefore have the potential to become the next generation of antibiotics. Some hybrid AMPs have been shown to possess improved activity (Shang et al., 2020; Wade et al., 2018, 2019) and here we explored the combination of two different classes of peptides, proline-rich peptides, and arginine-isoleucine-rich peptides. While the mode of action of the proline-rich peptides is known, the mode of action of the arginine-isoleucine-rich peptides has yet to be discovered. We hypothesized a “dirty drug” model and performed TEM studies on one of the arginine-isoleucine-rich peptides (optP7). The peptides induced outer membrane stress and created an enlarged and spherical nucleoid with some small denser foci, typically seen in DNA-dependent RNA polymerase inhibitors and protein synthesis inhibitors. In addition, a non-lytic mode of action was observed.

To discuss the obtained results in the context of conventional antibiotics, compared the peptides to three antibiotics, tigecycline (bacteriostatic activity by binding to the 30S ribosomal subunit of the bacterial ribosome), chloramphenicol (bacteriostatic activity by binding to the 30S ribosomal subunit of the bacterial ribosome), and rifampicin (bactericidal activity by inhibiting the DNA-dependent RNA polymerase activity), see Figure 5. This data was measured at the same beamtime session as the peptide set. As the antibiotics have slower kill kinetics than the AMPs, 240min incubation time was used. The concentration of the standard antibiotics (tigecycline, MIC 4µg/ml, chloramphenicol MIC 8µg/ml, rifampicin MIC 15µg/ml) was 3x their MIC against *E. coli*. The PCA showed that all peptides induced ultrastructural changes were distinctively different from conventional antibiotics. The PCA values of Bac5-v291 and hyP7B5GK deviate strongly from the untreated controls and the standard antibiotics. The PCA values of peptides hyP7B5Cys and optP7 are nearest to chloramphenicol.

**Figure 5:**
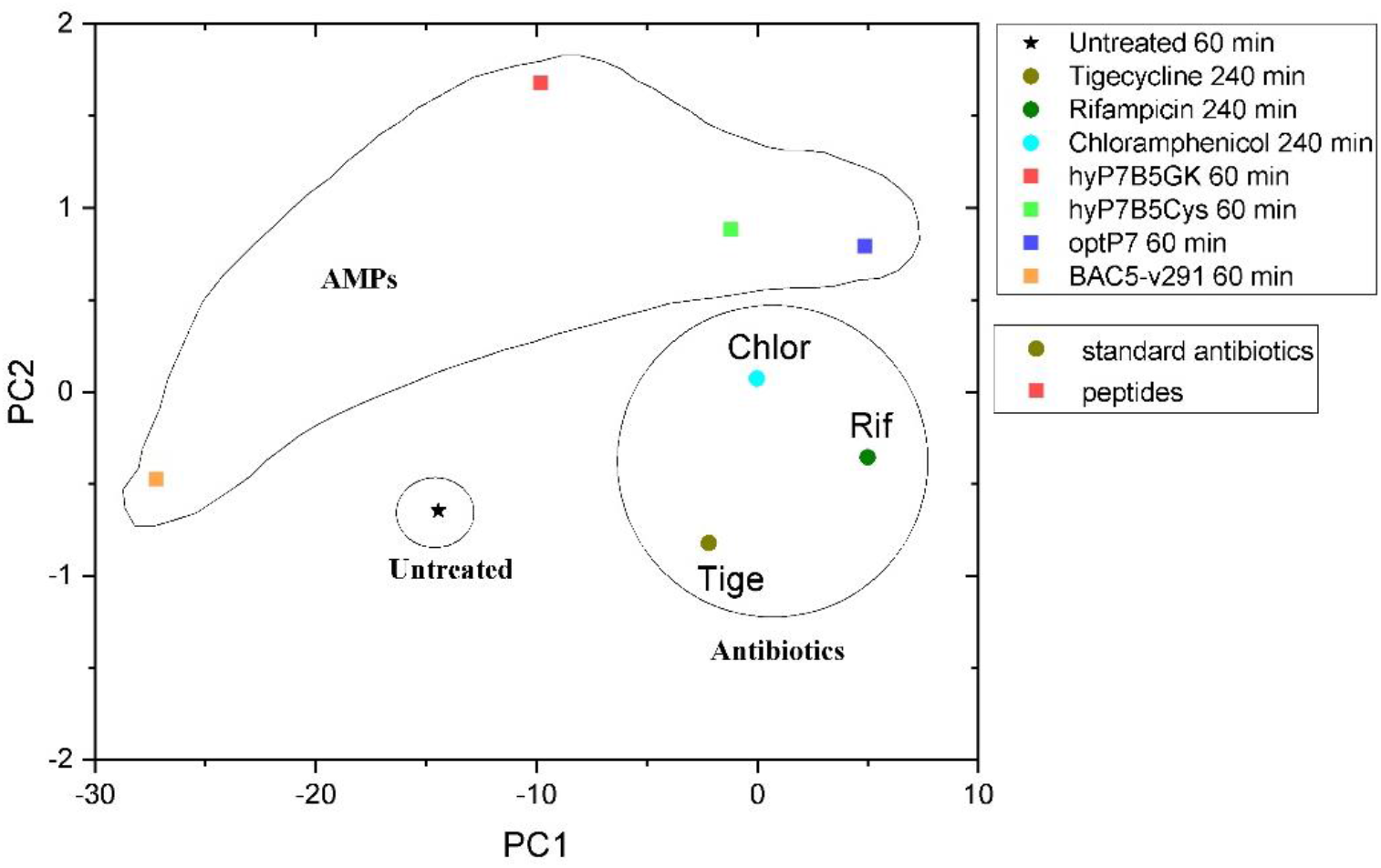
Comparison between principal component analysis (PCA) of the BioSAXS curves of *E. coli* treated with different AMPs and antibiotics.

Comparison between TEM pictures of *E. coli* treated with optP7, chloramphenicol, and tetracycline are shown in Figure 6. Very similar to the PCA of the BioSAXS data, the changes in *E. coli* treated with opP7 are distinctively different from the changes caused by the two antibiotics. They also show similarities, especially the enlarged and central nucleoid, and therefore explains well the closeness seen in the PCA. While TEM can only provide “snapshots” of all bacteria, BioSAXS reveals internal morphological changes averaged across hundreds of millions of bacteria, thus providing an unbiased representation of the treatment effects.

**Figure 6:**
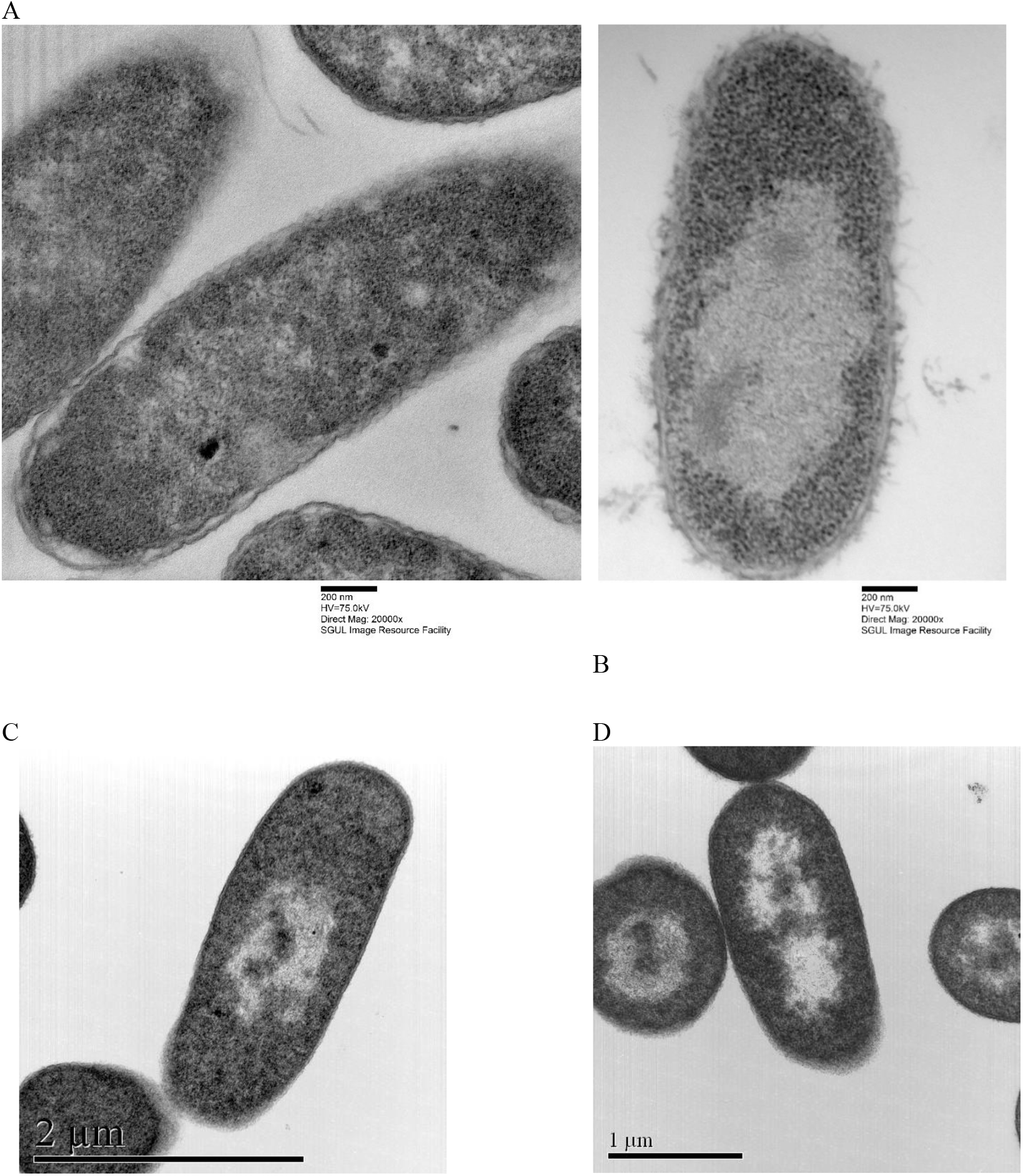
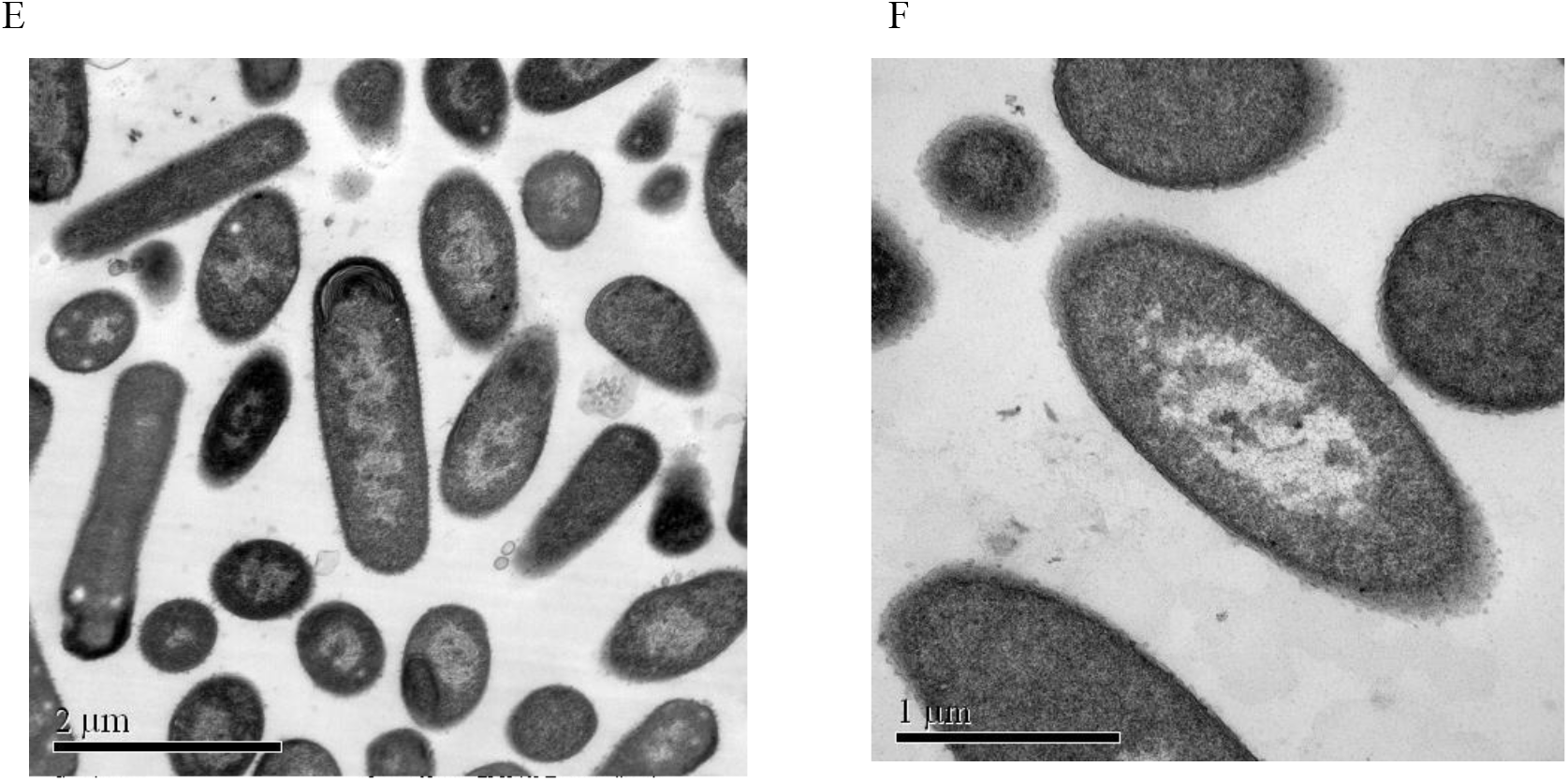
Transmission electron micrographs of *Escherichia coli*. A) Untreated samples, B) treated with optP7 for 60 min, at 32 µg/ml, C) and D) treated with chloramphenicol for 240 min, 60 µg/mL, E) and F) treated with tetracycline for 240 min, 30µg/ml.

All hybrid peptides showed a stronger activity against *E. coli* compared to the proline-rich ones, with Bac5-v291 as the only exception. The increase in activity compared to the arginine-isoleucine-rich peptides was up to 6-fold and dependent on the combination of peptides and bacteria. We have observed a very similar pattern while studying lipidation, glycosylation, cyclisation, and grafting into a cyclotide (Grimsey et al., 2020; Koehbach et al., 2021). All these modifications did not lead to a general improvement and were therefore not a “one fits all strategy”. A very important positive result was the high HC50 values of all peptides. Some of them showed a larger than 100fold therapeutic window (HC50/MIC). It demonstrates the potential of such compounds to be developed further into drug candidates against multi-drug resistant bacteria.

Interestingly, the hybrid peptide hyP7B5GK shows very different ultrastructural changes as both parents, indicating that both modes of action are creating ultrastructural changes at the same time and/or a novel mode of action occurred. In contrast, the hybrid molecule that is designed to fall apart in the bacterial cytosol by the reduction potential of that environment is closer to the parental peptide optP7, indicating a dominant contribution of the optP7 compound.

In conclusion, the creation of hybrid peptides leads in some cases to compounds with improved activity, which depends also on the target bacteria. Both parent peptides show unique effects on the ultrastructure of *E. coli* and the hybrid peptide hyP7B5GK shows yet another pattern of ultrastructural change, as measured by BioSAXS.

## 3 Author Contributions

K.H., conceptualization, resources (arginine-isoleucine-rich peptides, assay development), data analysis, supervision, funding acquisition, writing—original draft preparation, writing—review and editing; J.G., investigation, data curation (peptide synthesis and purification, MICs, haemolytic assay, TEM pictures); C.R. investigation, data curation (MICs, BioSAXS data analysis); N.S., investigation, data curation (MICs, BioSAXS sample preparation); P.M.L-P; investigation, data curation (peptide synthesis and purification); V.M.G., resources (BioSAXS); A.R.vG, investigation, data curation (TEM pictures); P.M., resource (BioSAXS), M.S., resource (proline-rich peptides), writing—review and editing; R.M., resource (BioSAXS data analysis), funding acquisition; A.R., resources (BioSAXS), supervision, funding acquisition, writing—review and editing. All authors have read and agreed to the published version of the manuscript.

## 5 Acknowledgements

We acknowledge the use of the Image Resource Facility, St George’s University. We thank Thomas Bruckdorfer from Iris Biotechnology for the discussion and recommendations of creating a peptide that will be cleaved in the cytosol. We acknowledge the team of P12 BioSAXS beamline at EMBL, Hamburg for excellent support. KH thanks the universe/life for the opportunity to continue on and despite the odds to be able to keep researching.

